# A Novel Bioprocess Control Strategy Under Uncertainty via Operational Space Identification

**DOI:** 10.64898/2026.02.07.704591

**Authors:** Mengjia Zhu, Oliver Pennington, Tararag Pincam, Mohammadamin Zarei, Sam Kay, Yongqiang Liu, Michael Short, Dongda Zhang

## Abstract

Bioprocesses are critical for sustainable industrial development but face challenges from their inherent uncertainties that affect efficiency and scalability. This study introduces a worst-case operational space design framework, integrating symbolic optimization with scenario-based validation, to preemptively mitigate uncertainties so that key performance indicators are consistently satisfied even under adverse conditions. The methodology is demonstrated through two case studies: a lab-scale fermentation for astaxanthin production and a site-scale anaerobic digestion for biogas production. Process uncertainty is quantified through model parameter variations (2% - 13%) to reflect real-world scenarios. The results indicate that highly flexible operational spaces are identified in both case studies, with control variables able to vary by up to 25% and 15% in the respective systems, while consistently adhering to system constraints. Validation over 10,000 scenarios confirmed 0 violation per case study. The proposed framework delivers validated operational flexibility with modest computational requirements, making it practical for lab-to-site deployment.

## 1 Introduction

Bioprocesses are important to a wide range of industries, including pharmaceuticals [1], biofuels [2, 3], food production [4], and environmental management [5, 6]. These processes rely on biological systems, such as cells, enzymes, or microorganisms, to convert raw materials into valuable products. Bioprocesses are typically characterized by their high specificity, energy efficiency, and ability to utilize renewable resources. Despite numerous advantages, bioprocesses face several challenges that impact their reliability, scalability, and efficiency [7], demanding precise control and monitoring to ensure consistent performance.

Traditionally, system identification techniques (e.g., dynamic modeling) are employed to develop mathematical models that describe the dynamic behavior of the bioprocesses [8]. These models are then used to design control strategies to maintain process variables within a desired range. For instance, Ramaswamy et al. [9] identified a nonlinear state-space model for a continuous stirred-tank bioreactor and used it to design an model predictive control (MPC) scheme that stabilizes an otherwise unstable operating point. Rogers et al. [10] uses transfer learning within a hybrid grey-box model to predict and optimize feeding strategies in high-cell-density microalgal fermentation. However, model uncertainties that stem from factors like incomplete knowledge of biological mechanisms, simplifications applied during model structure identification, inherent stochasticity in biological systems, and measurement errors pose challenges on the model construction and controller design [11, 12], leading to model-process mismatch, suboptimal performance, reduced yields, and increased costs, particularly in large-scale operations where deviations are magnified. Addressing these uncertainties is critical to improving bioprocess efficiency and scalability while maintaining the quality of the final product.

Significant efforts have been devoted to bioprocess modeling and optimization to effectively address uncertainty and enhance performance. Specifically, dynamic modeling and optimization approaches use physics-informed, data-driven, or hybrid models to predict and optimize dynamic systems effectively. Physics-based models utilize fundamental biological and chemical principles to construct mechanistic representations of the system, ensuring predictability for systems where the underlying processes are well understood. Conversely, data-driven models, such as artificial neural networks, bypass the need for extensive mechanistic knowledge by relying on experimental data to capture system behavior. For example, Del Rio-Chanona et al. [13] compared a mechanistic kinetic model against a data-driven artificial neural networks (ANN) for fed-batch microalgal lutein production and showed both improved intracellular lutein, with the ANN yielding higher total lutein and more accurate open-loop optimization, highlighting when data-driven surrogates can outperform kinetics in complex systems. Bradford et al. [14] developed a Gaussian-process–based dynamic model with uncertainty quantification for microalgal growth/lutein and carried out dynamic optimization under uncertainty, demonstrating Gaussian process (GP) models can match ANN predictive accuracy while providing principled uncertainty estimates for safer optimization.

On the other hand, hybrid models, which combine the strengths of physics-based and data-driven approaches, further enhance process optimization by integrating mechanistic insights with data-driven adaptability, enabling robust predictions across diverse operational conditions. For example, Zhang et al. [15] proposed a hybrid framework that filters noisy measurements with a simple kinetic model, trains a data-driven predictor, and re-fits the kinetic model as a soft sensor, enabling online monitoring and near-optimal fed-batch lutein production. Rogers et al. [16] systematically vary hybrid-model “greyness” by embedding different levels of kinetic structure and a GP component to learn unknown kinetics during fermentation of Cunninghamella echinulata, where they found too-strong mechanistic bias can hurt performance, while hybrids improve predictive confidence and scale-up temperature-shift predictions.

Several approaches have been developed to address model parameter uncertainties in bioprocess optimization and control. Among them, robust optimization (RO) draws theoretical interest due to its ability to provide reliable and stable solutions. RO aims to minimize the impact of process variability by ensuring that the chosen solution remains feasible and near-optimal even under the worst-case or most likely variations in model parameters [17, 18]. This is achieved by incorporating uncertainty explicitly into the optimization problem through deterministic equivalents or uncertainty sets, which define the bounds of parameter variability. For example, Santos et al. [19] formulate a min–max robust nonlinear MPC for fed-batch E. coli cultures with overflow metabolism, optimizing substrate feed while considering bounded uncertainties for 3 kinetic parameters. The scheme maintains performance under plant-model mismatch. Although RO methods provide guarantees over the uncertainty set, its implementation can be computationally demanding, especially for high-dimensional or nonlinear systems. Also, RO often results in conservative solutions and may be limited to specific types of uncertainties and problem structures [20, 21].

As an alternative to RO, scenario optimization assesses the impact of parameter variability by approximating the infinite set of uncertainties by a finite, representative set of scenarios generated through sampling techniques such as Monte Carlo (MC) or Latin Hypercube Sampling (LHS) [22, 23, 24]. Constraints are enforced for each scenario, transforming the problem into a finite but larger optimization problem. For example, Lucia and Engell [25] apply scenario tree to represent the uncertainty evolution for a batch bioreactor, taking future uncertainty realizations into account to improve constraint satisfaction and closedloop robustness. While this method improves computational tractability, it may require a large number of scenarios to achieve desired robustness, which can make their application prohibitive for complex or large-scale systems [26].

Beyond robust and scenario approaches, many workflows reduce computational cost via surrogate-based (meta-model) frameworks that emulate bioprocess models and thereby enable tractable uncertainty propagation and optimization under uncertainty [27]. Common choices include GP/Kriging surrogates (with predictive uncertainty that supports Bayesian, quantile- or reliability-based objectives) [28], polynomial response surfaces and (sparse) polynomial chaos expansions (PCE) [29], and multi-fidelity variants that blend cheap and high-fidelity simulations [30].

Within this surrogate family, PCE offers an analytic, spectral representation of input uncertainty with attractive properties for moment evaluation and sensitivity analysis. PCE represents uncertain model parameters as expansions in terms of orthogonal polynomials [31, 29], which transforms the stochastic nature of system inputs into a deterministic problem by approximating output uncertainties as a finite series of polynomial terms. Then, standard optimal control strategies can be applied to the deterministic model to determine control actions. By reducing the dimensionality of stochastic problems, PCE allows for the analysis of complex systems with minimal computational overhead while preserving accuracy, providing a computationally efficient alternative to more resource-intensive methods like MC simulations. For example, Paulson et al. [32] introduce a non-smooth PCE (nsPCE) tailored to dynamic flux balance analysis (dFBA), where they model the singularity (activeset switch) time to partition the parameter space and fit sparse, basis-adaptive PCEs on each smooth region, achieving over 800-fold speedups for uncertainty quantification (UQ) and Bayesian parameter estimation on an E. coli genome-scale dFBA with diauxic growth while maintaining accuracy. PCE is particularly effective for systems with well-defined probability distributions. Despite its advantages, PCE has limitations, including reduced accuracy for highly nonlinear systems or when input distributions deviate from assumed forms.

On the other hand, real-time feedback control (RTFC) strategies can be applied to adjust system behavior. RTFC uses advanced monitoring systems and adaptive feedback algorithms to dynamically adjust control actions in response to process deviations [33, 34, 35]. By continuously monitoring key process metrics, this approach ensures that the system remains within desired performance thresholds, even under fluctuating conditions. RTFC can enhance reliability and robustness but comes with high costs associated with infrastructure, such as sensors, actuators, and computational platforms [36, 37]. Consequently, its adoption is often limited to large-scale operations or industries with significant capital investment, while small-to medium-scale facilities may find it financially unfeasible.

Despite advancements in bioprocess modeling and optimization under uncertainty, relatively little effort has been directed toward developing innovative control methods [38, 36]. Many biochemical and formulated chemical industries face challenges in strictly adhering to a fixed set-point control trajectory due to various factors, such as human error, system inconsistencies, and the difficulty of measuring key states in real time due to the inherent complexity of bioprocesses, which can lead to unnecessary energy consumption. To address this challenge, we propose a flexible bioprocess control strategy via operational space identification for model uncertainty management. Specifically, in this paper, we introduce a worst-case operational space design (WC-OSD) framework to identify a flexible operational space for control actions, such that, if we operate within this space, key performance indicators (KPIs) can be achieved, even under adverse conditions. By shifting from a rigid set-point-based approach to an operational space-based paradigm, this framework enhances both flexibility and efficiency while reducing dependence on frequent monitoring and control adjustments. Additionally, by optimizing operational flexibility in advance—based on system models and identified uncertainties derived from model construction or historical data—WC-OSD provides a scalable solution applicable to various bioprocesses, from small-scale facilities to large industrial operations, making it a more attractive alternative for real-world applications.

Key features of the WC-OSD framework include: 1) explicit incorporation of process uncertainties into operational planning, where these uncertainties are quantified through variations in model parameters; 2) integration of symbolic frameworks in optimization to preserve model fidelity and ensure constraint satisfaction; and 3) implementation of a scenario-based validation procedure to enhance the reliability of the identified operational space.

The rest of the paper is organized as follows. The target problem is described in Section 2, with the proposed solution methods detailed in Section 3. The effectiveness of the proposed approach is demonstrated via two case studies—one on a lab-scale fermentation process of astaxanthin production, and another on a site-scale anaerobic digestion (AD) reactor—discussed in Section 4. Lastly, conclusions and future work are noted in Section 5.

## 2 Problem Description

Consider a bioprocess described by a mechanistic model in the form of ordinary differential equations (ODEs) in Equation (1):

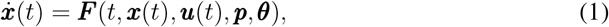

where, ***x*** is the vector of state variables (*e*.*g*., biomass concentration, substrate concentration), and 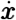 is the corresponding time derivative. ***u*** is the vector of control inputs (*e*.*g*., feed rate, temperature). And ***p*** represents parameters subject to bounded uncertainties, *i*.*e*., ***p*** ∈ [***p***_min_, ***p***_max_], with ***θ*** being the vector of fixed parameters.

The following assumptions are made in this study: (*i*) the process uncertainties are quantified via uncertain model parameters, whose range of variations is known. The parameter ranges can be taken from reported experimental confidence intervals or be determined from parameter estimation during model construction; (*ii*) the initial state of the system is known; (*iii*) the desired output (i.e., KPI) and its relevant acceptable ranges have been defined; (*iv*) the control input can be time-varying, and a reference trajectory is provided (e.g., based on the historical data) or can be determined via optimal control strategies; (*v*) the confidence band to be identified for the control input can be time-varying, and the system operates on a fixed time horizon; and (*vi*) the system dynamics can be described by an adequate model, which is provided or can be constructed.

The goal is to identify a practical and flexible operational space for ***u***(*t*) such that, if we operate within this space, the desired KPIs of the bioprocess are likely to be consistently met, independent of the uncertain model parameters.

## 3 Methodology

### 3.1 Optimization problem formulation

The operational space identification problem is translated into an optimization problem. We first discretize the time horizon [0, *T* ] into *N* intervals with piecewise constant controls ***u***_*k*_ for *k* = 0, 1, …, *N* − 1. The discretized time steps *t*_*k*_ = *k*Δ*t*, for *k* = 0, 1, …, *N*, where 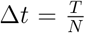. Our objective is to find the largest operational space for the control inputs subject to process constraints. Specifically, we consider maximizing the difference between the upper 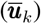 and lower 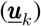 bounds of the control inputs over the time horizon described in optimization problem (2):

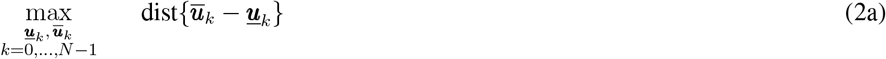

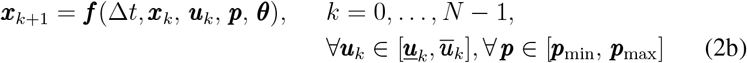

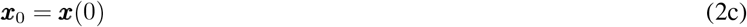

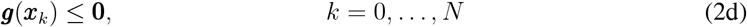

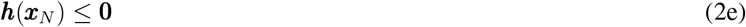

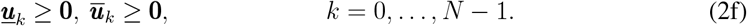

Here, (2b) represents the discretized system dynamics with ***f*** (·) being the selected numerical integrator (*e*.*g*., implicit/explicit Euler, Runge-Kutta, *etc*.), (2c) denotes the initial condition of the system, (2d) and (2e) defines the path and terminal constraints (e.g. KPI specifications), respectively, and (2f) defines physical feasibility.

### 3.2 Solution approach

The proposed solution approach includes four main steps which are summarized as follows. The first step involves defining a nominal control trajectory, ***ũ***_*k*_, for discrete-time steps *k* = 0, …, *N* − 1. In the second step, an optimization-based approach is employed to determine the worst-case combination of uncertain model parameters. This identified combination is then used in the next step to define the operational space, ensuring that the system maintains reliable performance even under the most adverse uncertainty conditions. In the third step, another two optimization problems are formulated and solved to identify the lower and upper bounds of the control variables. These bounds collectively form the operational space. Finally, a scenario-based verification step is conducted, where the identified bounds are validated using an MC sampling. In the following, we provide details regarding each step.

#### 3.2.1 Nominal control trajectory identification

The nominal trajectory, ***ũ***_*k*_, serves as a baseline for subsequent analysis and can be determined in two main ways: it may either be predefined based on prior knowledge (e.g., historical operational data) or computed dynamically using an optimal control strategy. To generate the nominal control trajectory, a standard dynamic optimization problem is formulated. The objective is to optimize the control variables while assuming that the uncertain model parameters, ***p***, are at their nominal values. A common method for defining these nominal values is to take the midpoint of the uncertainty ranges, expressed as 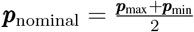, where ***p***_max_ and ***p***_min_ represent the upper and lower bounds of the uncertain model parameters, respectively. This simplification allows for a deterministic optimization problem that minimizes or maximizes a chosen performance criterion, such as energy consumption, tracking error, or system cost, subject to dynamic and operational constraints.

The general form of the optimization problem is given as:

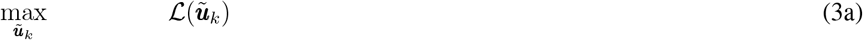

subject to:

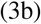

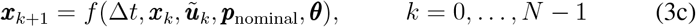

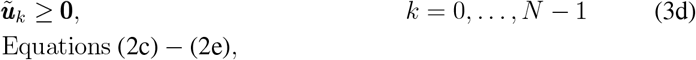

where ℒ (·) is the objective function. Note, maximizing ℒ (·) is equivalent to minimizing −ℒ (·). And the solution of (3) is optimal under nominal settings.

#### 3.2.2 Worst-case parameter combination identification

After determining the nominal control trajectory ***ũ***_*k*_, we investigate how uncertain model parameters could degrade system performance by identifying their worst-case combination under nominal control. To this end, we define a quantitative metric *J* (***p***) that captures undesirable behavior (e.g., large deviations from a target trajectory or other adverse outcomes) and explicitly depends on the uncertain model parameter vector ***p***. Using this metric, we formulate an optimization problem (see Problem (4)) to find the parameter values that produce the worst performance while respecting the system’s operational constraints. In Problem (4), we assume that the nominal control trajectory can still satisfy key performance indicators even under these worst-case parameter variations:

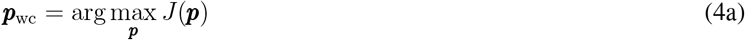

subject to:

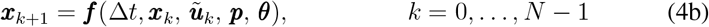

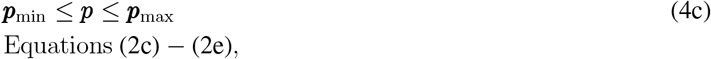

where ***p***_wc_ is the determined worst-case uncertain model parameter combinations. By using the optimization approach, it is possible to identify the worst-case combinations that do not occur at the boundaries. Nevertheless, it is important to note that the determined worst case ***p***_wc_ is conditioned on the given nominal control trajectory ***ũ***_*k*_. Therefore, subsequent validation step is necessary when the operational space is established based on ***p***_wc_.

#### 3.2.3 Worst-case-based operational space design

With the identified ***ũ***_*k*_ and ***p***_wc_, we determine the lower and upper bounds for the control inputs around ***ũ***_*k*_ independently by maximizing the distance between the two whilst satisfying process constraints. The lower bounds are identified using the optimization problem (5), which is formulated to maximize the minimum weighted difference between the lower bounds, ***<u>u</u>***_*k*_, and the nominal values, ***ũ***_*k*_:

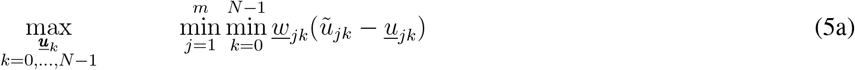

subject to:

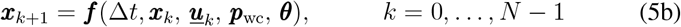

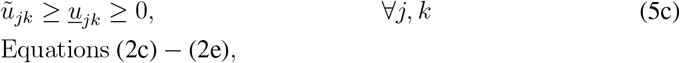

where *u*_*jk*_ is the lower bound for the *j*-th control input at time step *k*, and *<u>w</u>*_*j,k*_ normalizes each control input’s contribution at that time step. Here, *j* indexes the control inputs ***u***_*k*_, and *m* is the total number of control variables in the bioprocess. This *max-min* formulation is preferred over maximizing the sum (*max-sum*) of differences to avoid situations where only a few control inputs exhibit significant deviations from the nominal values, while most deviations remain near zero. Such imbalanced adjustments can lead to less practical bounds, potentially resulting in infeasible or inefficient control actions. These arguments will be further illustrated in the first case study (see Section 4.1), where a comparison study between *max-min* formulation and *max-sum* is performed.

Similar to the lower bounds identification, the upper bounds 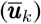 can be identified by solving the following optimization problem, with 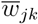 being the standardization parameters:

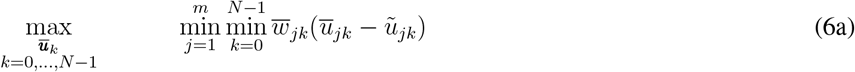

subject to:

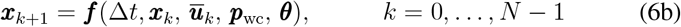

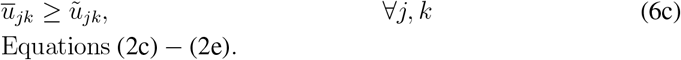

It is important to note that the formulations are flexible in handling restrictions on the variability of control inputs. For instance, actuator or operational constraints such as rate-of-change limits, maximum allowable deviations, or bounds on the number of adjustments can be included as additional constraints. If partial violations are acceptable in practice, slack variables or soft constraints can be introduced and penalized in the objective function.

#### 3.2.4 Operational space validation

Since the worst-case parameter combination is identified based on nominal control actions, it is important to validate the effectiveness and practicability of the identified operational space against a variety of conditions. Here, we apply MC simulation. First, a large number of uncertain model parameter combinations are randomly sampled from their uncertainty bounds. Each combination will then be coupled with a control trajectory randomly sampled from the defined operational space, forming one scenario of the system. Every scenario is simulated via the system model and the system’s responses are evaluated against KPIs and other system constraints. The outcomes are then analyzed to determine if there exists any instance that fails to meet the requirements. If the required robustness (e.g., % violation ≤ *ξ*, where *ξ* is a pre-defined threshold) is not met, we can narrow the current operational bounds by feeding the identified violation uncertain model parameter combinations into the optimization problems (7) and (8):

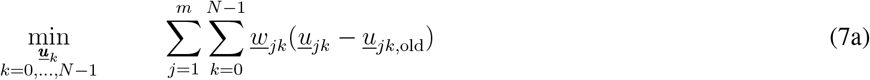

subject to:

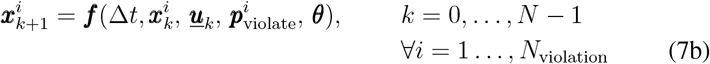

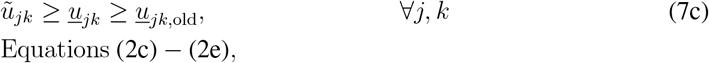

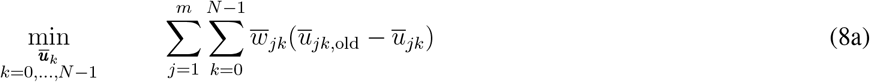

subject to:

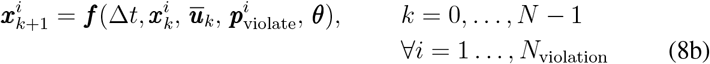

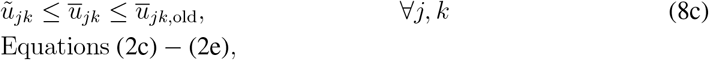

where the previously identified lower and upper bounds were assigned to *u*_*jk*,old_ and 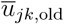, with *i* denotes the index of uncertain model parameter combination, 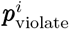, that violates the constraints and *N*_violation_ being the total number of violated cases identified during MC simulation. In this refinement step, different from optimization problems (5) and (6), we identify the new bounds by minimizing the total weighted decrease from the previous bounds to the new bounds, making only necessary reductions to satisfy the accumulated constraints introduced by 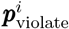 . This new formulation avoids the necessity to re-run previously sampled uncertain model parameter combinations, therefore reducing the computational costs. The updated bounds are then validated with a new MC simulation. We repeat the cycle of MC simulation and re-optimization with (7) and (8) until the required robustness level is achieved. Unless stated otherwise, uncertain model parameter combinations are sampled uniformly over their bounds. If reliable parameter distributions are available, samples may instead draw from those distributions to estimate distribution-aware violation rates.

The WC-OSD framework described in this section is summarized in Figure 1. WC-OSD provides a structured approach to managing uncertainty in bioprocess control by systematically identifying feasible operational spaces. By integrating optimization with worst-case scenario analysis, this framework prompts reliability of the control strategy without excessive reliance on real-time adjustments. The final validation step further strengthens the approach by verifying performance across a range of variations.

**Figure 1.**
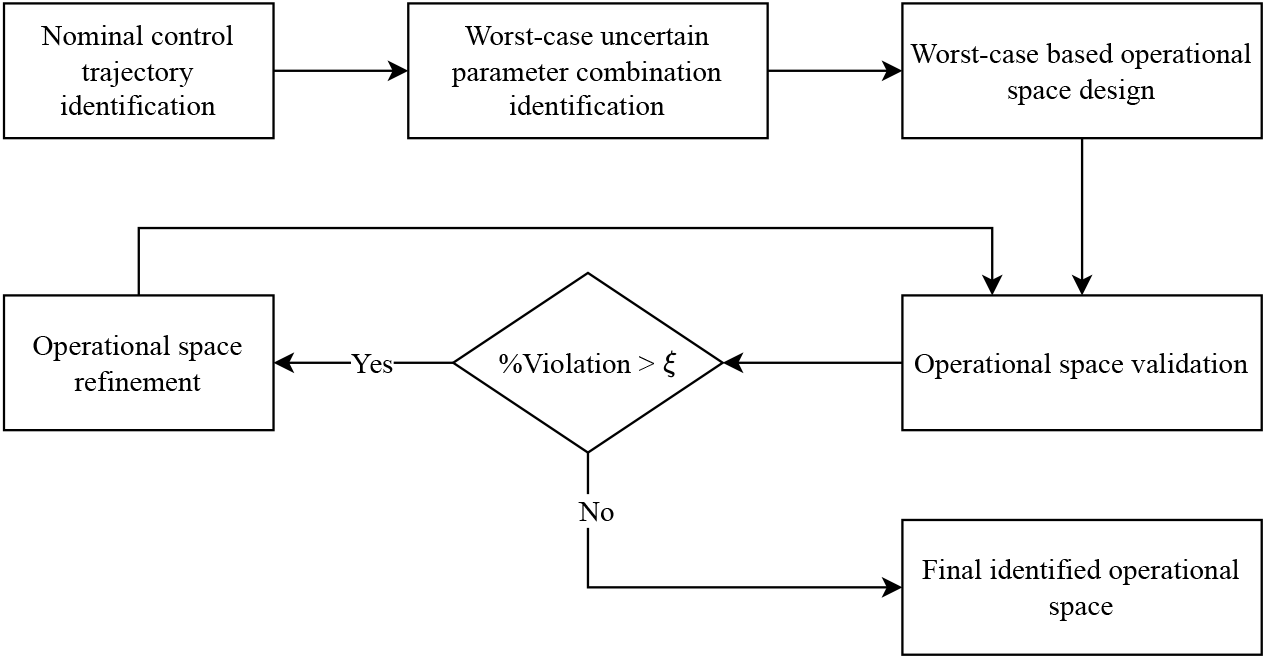
Flowchart of the WC-OSD framework.

## 4 Case Studies

The proposed WC-OSD approach described in Section 3 is applied to two case studies: one on a lab-scale fermentation process of astaxanthin production, and another on a site-scale anaerobic digestion (AD) reactor. For the first case study, the system model is taken from the literature, while for the second case study, the model is constructed based on four lab-scale digesters and one site-scale digester, being fed the same feedstock combinations.

### 4.1 Fermentation process for astaxanthin production

Astaxanthin, a commercially valuable carotenoid, plays an important role across industries, from natural red colorants in food and cosmetics to powerful antioxidants in nutraceuticals and medicine [39]. Despite its high market value, existing production methods, such as microalgal biosynthesis, face engineering challenges including low growth rates and the complexity of scaling up photobioreactors. As a result, yeast fermentation, particularly using Xanthophyllomyces dendrorhous, has been considered as the major approach for large scale astaxanthin production [40]. Recent studies emphasize the potential of mixed sugar fermentation, utilizing sugars derived from biowaste, as a sustainable and economically viable approach. Vega-Ramon et al. [41] proposed a kinetic model of this fermentation process to model the effects of sugars on yeast biomass growth and astaxanthin production, which is converted to a fed-batch model by including the feed flow rate (g/L), *F*_in_(*t*), as the control input. The resulting model is presented in Equation (9):

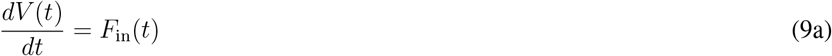

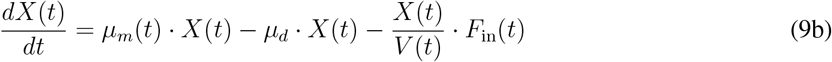

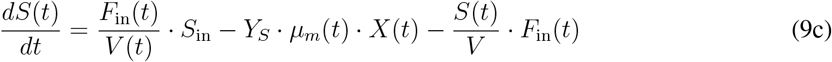

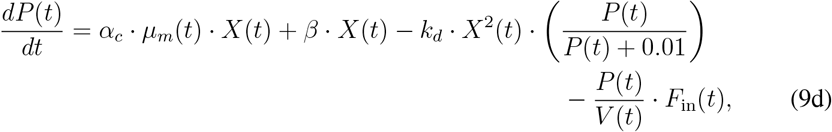

Where 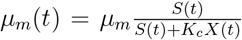. Here, *X*(*t*) is the biomass concentration (g/L), *S*(*t*) is the substrate concentration (g/L), *P* (*t*) is the product concentration (mg/L), and *t* is time (h). *S*_in_ is the inflow substrate concentration, which is assumed to be constant. The parameters *µ*_*m*_ and *µ*_*d*_ represent the maximum specific growth rate and the specific decay rate, respectively. *Y*_*S*_ is the substrate yield coefficient, with *α*_*c*_ and *β* being the growth-dependent yield coefficient and growth-independent yield coefficient, respectively. *k*_*d*_ is the specific consumption rate, and *K*_*c*_ is the substrate saturation constant. The value of each parameter is shown in Table 1.

**Table 1:** Model parameters for astaxanthin production [41].

| Parameter | Nominal Value |
| --- | --- |
| $\mu_m$ (h<sup>-1</sup>) | 0.43 |
| $K_c$ ([-]) | 63.7 |
| $\mu_d$ (h<sup>-1</sup>) | $2.1 \times 10^{-3}$ |
| $Y_S$ (g/g) | 2.58 |
| $\alpha_c$ (mg/g) | 7.88 |
| $\beta$ (mg/g h<sup>-1</sup>) | 0.236 |
| $k_d$ (mg L g<sup>-2</sup> h<sup>-1</sup>) | 0.0648 |

Let ***x***(*t*) = [*V* (*t*), *X*(*t*), *S*(*t*), *P* (*t*)]^⊤^, *u*(*t*) = *F*_in_(*t*), ***p*** = [*µ*_*m*_, *K*_*c*_, *µ*_*d*_, *Y*_*S*_, *α*_*c*_, *β, k*_*d*_]^⊤^, and *θ* = *S*_in_. The system can be compactly written as:

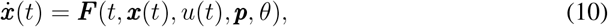

which is consistent with the general problem formulation (1).

#### 4.1.1 Operational space design

For illustrative purpose of the proposed method, we consider uncertainties in the parameters with variations of ±2% around their nominal values (Table 1). In applications, the variation ranges can be set from parameter-estimation posteriors or confidence intervals. The initial volume (*V* (0)), biomass concentration (*X*(0)), substrate concentration (*S*(0)), and product concentration (*P* (0)) are 0.4 L, 0.1 g/L, 6 g/L, and 0 mg/L, respectively. The substrate concentration in the feed *S*_in_ is set to 12 g/L, and the reactor’s volume is 2 L (*V*_reactor_). The process is run for 168 hours (*T* ). The feed flow rate *F*_in_(*t*) is adjusted every 12 hours, leading to 14 discretized time segments (*N* ). The KPI in this case is to ensure that the mass production of astaxanthin at the end of the time horizon is higher than 58 mg, *i*.*e*., *V* (*N* )*P* (*N* ) ≥ 58 mg. Several path constraints are involved: (*i*) all the state variables are non-negative, *i*.*e*., ***x***(*t*) ≥ 0, for *t* ∈ [0, *T* ]; (*ii*) maximum working volume should not exceed 95% of the reactor’s volume, *i*.*e*., *V* (*t*) ≤ 0.95*V*_reactor_, with the nominal working volume being less than or equal to 80% of the reactor’s volume; (*iii*) no feed in the first and last days (24 hours), *i*.*e*., *u*_*k*_ = 0, for *k* ∈ {0, 1, *N* − 3, *N* − 2}; (*iv*) the substrate concentration should not change dramatically, *i*.*e*., 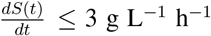 and (*v*) the mass of the product astaxanthin should increase throughout the process, *i*.*e*., 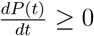.

To identify the nominal control trajectory, *ũ*_*k*_, we set the uncertain model parameters to their nominal values as in Table 1. The objective function ℒ (·) is formulated as maximizing the astaxanthin production, i.e., ℒ (*ũ*_*k*_) = *V* (*N* )*P* (*N* ), while satisfying all the aforementioned constraints. On the other hand, when determining the worst-case parameter combination, we set *J* (***p***) = −*V* (*N* )*P* (*N* ), i.e., finding the parameter combination that leads to the lowest astaxanthin production under nominal operating conditions.

As for the operational space identification, since it is not unique, additional constraints can be introduced to shape its profile. For example, if it is known in advance that control is critical at a certain time step and limited variation is preferred, constraints can be added to enforce it. In our case, we introduce constraints that increase control variability in the middle time segments over the time horizon, specifically, 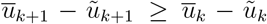 and *ũ*_*k*+1_ − *u*_*k*+1_ ≥ *ũ*_*k*_ − *u*_*k*_ for *k* ∈ {3, …, *N* − 5} . The rationale is that by closely following the desired operation in the early stages, we allow for greater flexibility in the later stages of the process.

Also, as noted in Section 3.2.3, *max-min* formulation is preferred over *max-sum* formulation when identifying the initial operational space. For comparison, we also solve the same operational space identification problem for the current case study via *max-sum* formulation of (5) and (6) (i.e., replace 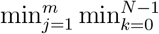 with 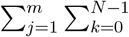 in both (5) and (6)).

All the computations are performed on an Intel i7-13700H 2.40-GHz CPU laptop with 32GB of RAM. To ensure feasibility with regarding the system dynamics and constraints when solving optimization problems involved in each step, a symbolic framework is incorporated into the optimization procedure. We use CasADi [42], a powerful tool tailored for handling optimal control problems. Specifically, direct transcription is used to transform (10) into the finite-dimensional nonlinear programming problem (NLP) in (2)-(8), with an explicit Runge-Kutta integrator for *f* (·). System dynamics, constraints, and objectives are built symbolically in CasADi to enable exact Jacobinas/Hessians via automatic differentiation, and solved with a sparse NLP solver IPOPT [43].

#### 4.2.4 Results and discussions

The worst-case parameter combinations, ***p***_wc_, was identified by solving (4), which took 376.1 seconds of CPU time, and was found to be [0.421, 65.0, 0.00214, 2.63, 7.72, 0.231, 0.0661]. The determined nominal control trajectory, *ũ*_*k*_, and the resulting operational spaces solved via *max-sum* and *max-min* formulations are shown in Figure 2. It took 0.2969 and 2.125 seconds of CPU time, respectively, to solve the *max-sum* and *max-min* formulations. It is observed that the *max-sum* approach (see Figure 2a) leads to a larger total deviation from the nominal control inputs (0.0459 L/h) than the *max-min* approach (0.0399 L/h) (see Figure 2b), but the deviations in Figure 2a are concentrated in only two segments. In contrast, the *max-min* approach distributes the deviations across all segments, allowing deviations of up to 25% from the nominal control trajectories. These findings highlight the effectiveness of the *max-min* formulation in identifying an operational space that is more practically advantageous.

**Figure 2.**
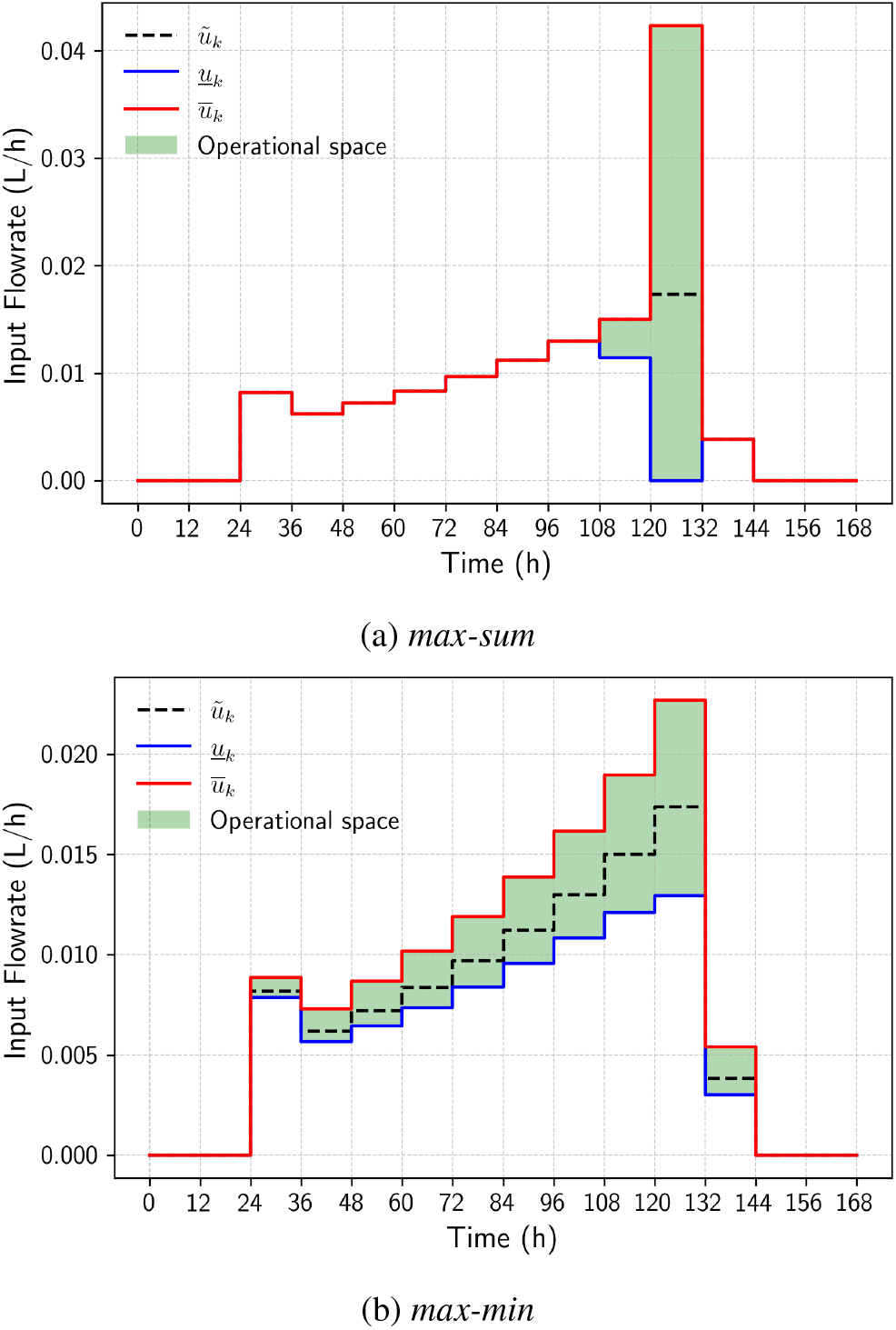
Comparison of *max-sum* and *max-min* optimization results for operational space identification of astaxanthin production

The operational space identified through the *max-min* formulation needs to satisfy all the path constraints and the terminal constraint that astaxanthin production is no less than 58 mg. Once the operational space is determined, it is validated via MC simulation, as described in Section 3.2.4, with a robustness threshold (i.e., the maximum allowed violation rate) *ξ* of 2%. Specifically, 10,000 random uncertain model parameter combinations are sampled within their bounds uniformly, each paired with a randomly generated control trajectory from the uniform distribution of the identified operational space. Figure 3a shows the mean and variation range of 10,000 control trajectories generated during MC validation, demonstrating a comprehensive coverage of the operational space depicted in Figure 2b. Additionally, the tested uncertain model parameter combinations and the corresponding KPI values are presented in Figure 3b. The parallel coordinate plot provides a detailed visualization of the sampled uncertain model parameter combinations, highlighting the thorough consideration of parameter variability during the validation process. In the plot, the left-most vertical axis represents KPI satisfaction, specifically whether the final astaxanthin mass (*V* (*N* )*P* (*N* )) exceeds 58 mg, while the subsequent vertical axes correspond to individual parameters and their respective uncertain ranges. The lines connecting these axes represent the uncertain model parameter combinations and the resulting mass production outcomes for each sampled scenario. Across all 10,000 sampled scenarios, no KPI violations were observed, showcasing the effectiveness of the operational space based control strategy.

**Figure 3.**
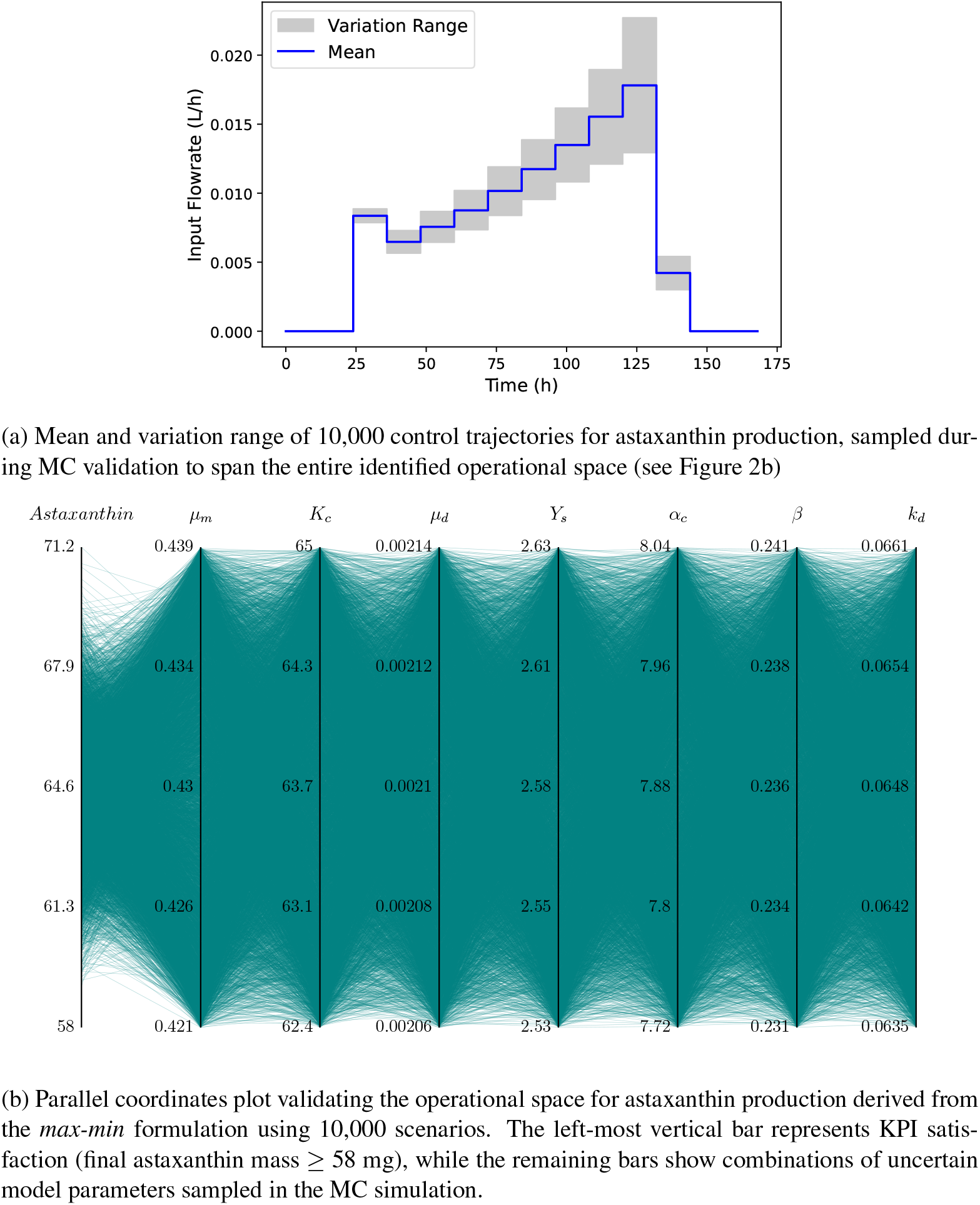
Operational space validation through MC simulation.

### 4.2 Anaerobic digestion for biogas production

For the second case study, we consider anaerobic digestion (AD) for biogas production under uncertainty. AD is an important technology in advancing sustainable energy systems. It transforms organic waste into biogas and nutrient-rich digestate, addressing critical challenges in energy transition and waste management [5, 6]. Biogas, a high-energy renewable fuel, can displace fossil fuels in heat and electricity, contributing to carbon neutrality goals [44]. Despite its potential, AD systems face challenges that undermine their efficiency and economic viability [7]. The biological complexity of AD introduces variability in methane yield, while feedstock heterogeneity and fluctuating operational conditions add layers of uncertainty. These challenges are further compounded by fluctuating market dynamics and the high costs associated with advanced real-time monitoring systems [45, 36, 37].

In this case study, we aim to design an operational space for the control variable (the hydraulic loading rate, *HLR*), such that if we operate within this space, the defined KPI (the biogas production) can be reliably achieved. We first discuss the AD model construction, parameter estimation, and validation on 4 lab-scale digesters and 1 site-scale digester. Note, the 4 lab-scale digesters were operated to mimic the conditions of the site-scale digester. Then, the nominal operating conditions are selected, based on which, the operational space is obtained using the proposed approach.

#### 4.2.1 Model construction

AD utilizes a variety of feedstock to generate biogas. Since feedstock is added every 1-2 hours and the digester retention time generally exceeds 30 days, the feeding can be approximated as continuous. Operating with constant volume and assuming Monod kinetics, a proposed mass balance model for the continuous system can be given by:

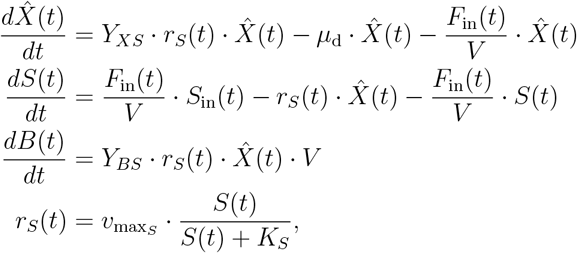

where *V* is the working volume in L, *F*_in_ is the flow passing through the digester in L d^−1^, *X* is cell density in unit dry weight per liter (uDW L^−1^), *S* is substrate concentration in gVS L^−1^ (note that VS stands for volatile solids), and *B* is biogas volume in L at standard pressure and temperature (note that the data was collected at specific pressure and temperature, which was later converted to standard conditions). The parameters 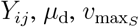, and *K*_*S*_ represent the yield of production of *i* with respect to component *j*, the specific cell death rate, the maximum specific substrate consumption rate, and the substrate concentration for which half the maximum rate is achieved, respectively.

Substrate intake concentration is approximated as 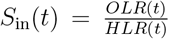 where *OLR*(*t*) is the organic loading rate in gVS L^−1^ d^−1^, and *HLR*(*t*) is the hydraulic loading rate in L_feed_ L^−1^ d^−1^, which gives 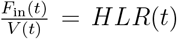. This means the final model can be written as in Equation (11):

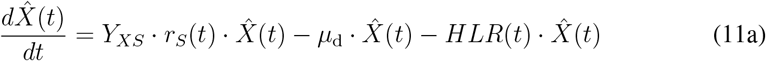

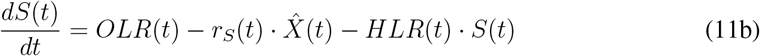

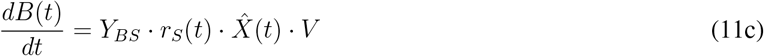

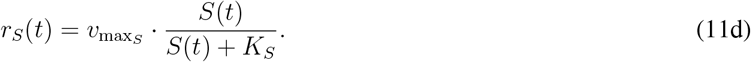

It should be noted that the biogas is not diluted by the hydraulic loading as it is emitted as a gas and removed from the top of the digester; 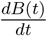 represents the rate of production of biogas in L d^−1^, not its rate of accumulation within the digester.

##### Kinetic model parameter estimation

The data available includes organic and hydraulic loading rates for each day, as well as daily biogas production, for both lab and site scales. In order to estimate the model parameters, an objective function was utilized to minimize the difference between the actual and simulated biogas production per day (Δ*B*) when using the recorded *OLR* and *HLR* inputs, as in Equation (12):

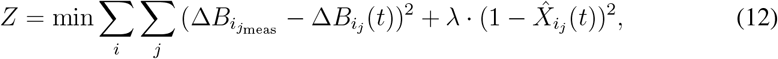

where *i* denotes the different digesters used for parameter estimation, and *j* denotes the specific data point for the given digester. During parameter estimation, a penalty term, with weight *λ*, was introduced to penalize changes in cell density during stable operation to identify a model that is able to mimic stable behavior. Increasing *λ* increases the fixing of cell density to unity during parameter estimation, however, this will result in decreased model fitting of the parameter estimation dataset. Due to the unavailability of cell density measurements, the model uses a standardized cell density, hence the penalty term pushing cell density to unity. The model parameters were estimated simultaneously on two lab-scale digesters and one real site, and validated on two other different lab-scale digesters. The lab-scale and real-site simulations share all parameters but were allowed to take different initial conditions. The real site data had one additional parameter (*α*) added to account for differences in production due to scale-up, as in Equation (13):

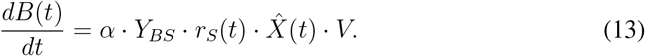

which replaces Equation (11c) for the site case.

The Mean Average Prediction Error (MAPE) beyond the initial settling period for the fitting of the lab-scale digesters to Equation (11) (using Equation (12)) is just 6.4%, with the real site counterpart having a MAPE of 8.5% when incorporating Equation (13). The parameter values are provided in Table 2, along with the considered uncertainty bounds which will be elaborated in Section 4.2.3 when discussing the operational space design. The fact that the parameter *K*_*S*_ is much larger than the simulated substrate concentration implies that the system is effectively operating under first-order kinetics with respect to substrate concentration. The parameter *α* being greater than unity implies that the site-scale digester produces a larger volume of biogas per unit of organic loading, which could be explained by the differences seen in biomass composition between the lab-scale and site-scale digesters, or even seasonal effects seen on substrate that can differ between the digesters. However, the fact that the parameter *α* is close to unity demonstrates the capability of the model to describe the dynamics of both lab-scale and site-scale digesters.

**Table 2:** Model parameters for ADM model described by Equations (11) and (13)

| Parameter | Nominal Value | Uncertainty |
| --- | --- | --- |
| $Y_{XS}$ (uDW gVS<sup>-1</sup>) | $2.63 \times 10^{-3}$ | $\pm 12\%$ |
| $\mu_d$ (d<sup>-1</sup>) | $4.89 \times 10^{-6}$ | $\pm 10\%$ |
| $Y_{BS}$ (L gVS<sup>-1</sup>) | $4.44 \times 10^{-4}$ | $\pm 13\%$ |
| $v_{\max_S}$ (gVS uDW<sup>-1</sup> d<sup>-1</sup>) | $4.76 \times 10^5$ | $\pm 5\%$ |
| $K_S$ (gVS L<sup>-1</sup>) | $5.08 \times 10^5$ | $\pm 2\%$ |
| $\alpha$ ([-]) | 1.35 | $\pm 6\%$ |

As can be seen in Figures 4 and 5, near-unity is maintained for cell density throughout the dynamic process for both the lab-scale and site-scale operations, as desired by the penalty term in Equation (12). It is also observable that substrate is predicted to maintain a concentration of similar value within digesters at both lab-scale and site-scale, averaging around 4-6 gVS L^−1^. This demonstrates reliability in the model as the lab-scale digesters were operated to imitate operations of the site-scale digester, so it is expected for state variables to be of comparable value.

**Figure 4.**
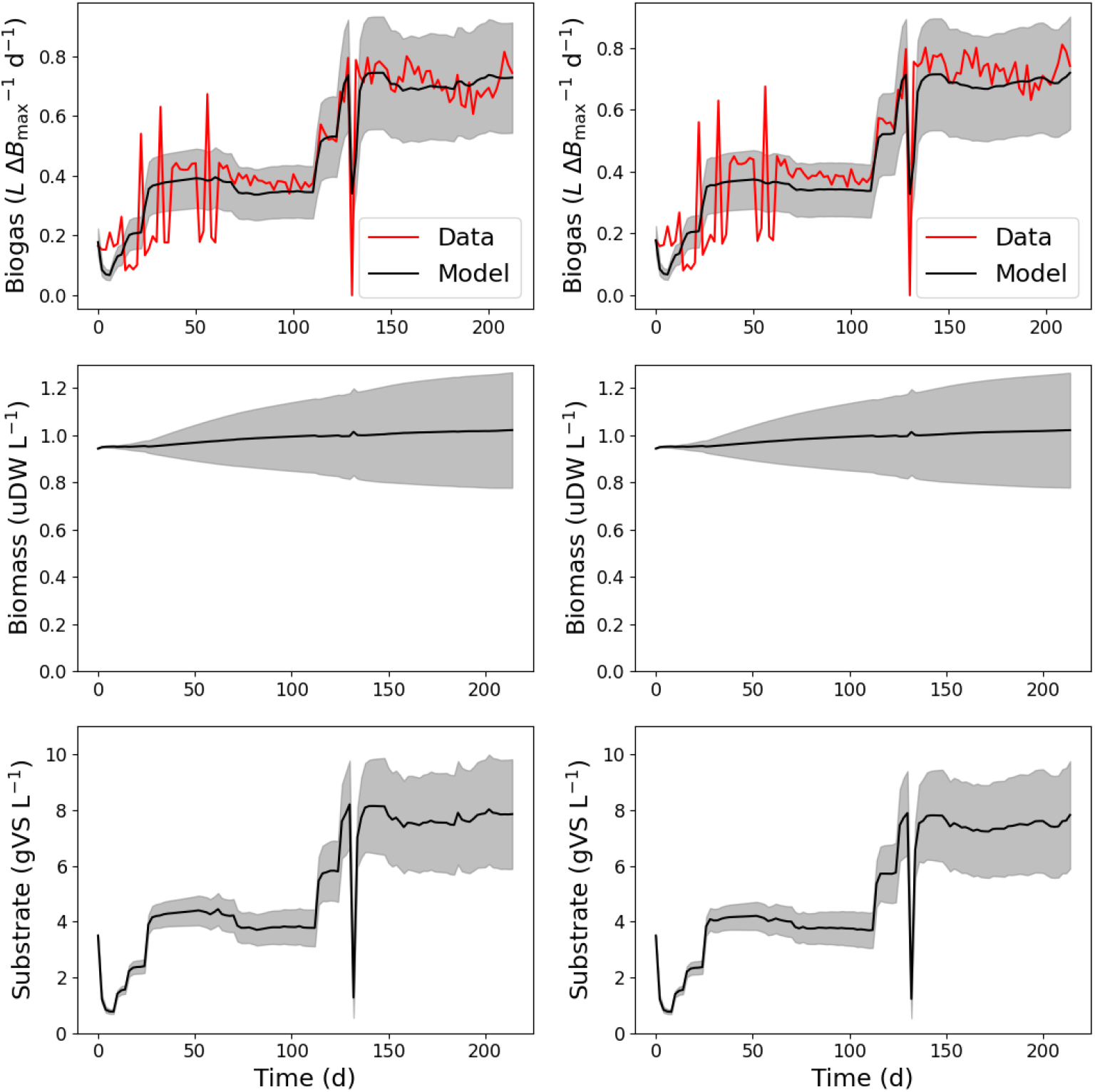
Parameter estimation fitting for lab-scale digesters 1 (left) and 2 (right). Gray bands are the model uncertainty (95% confidence interval)

**Figure 5.**
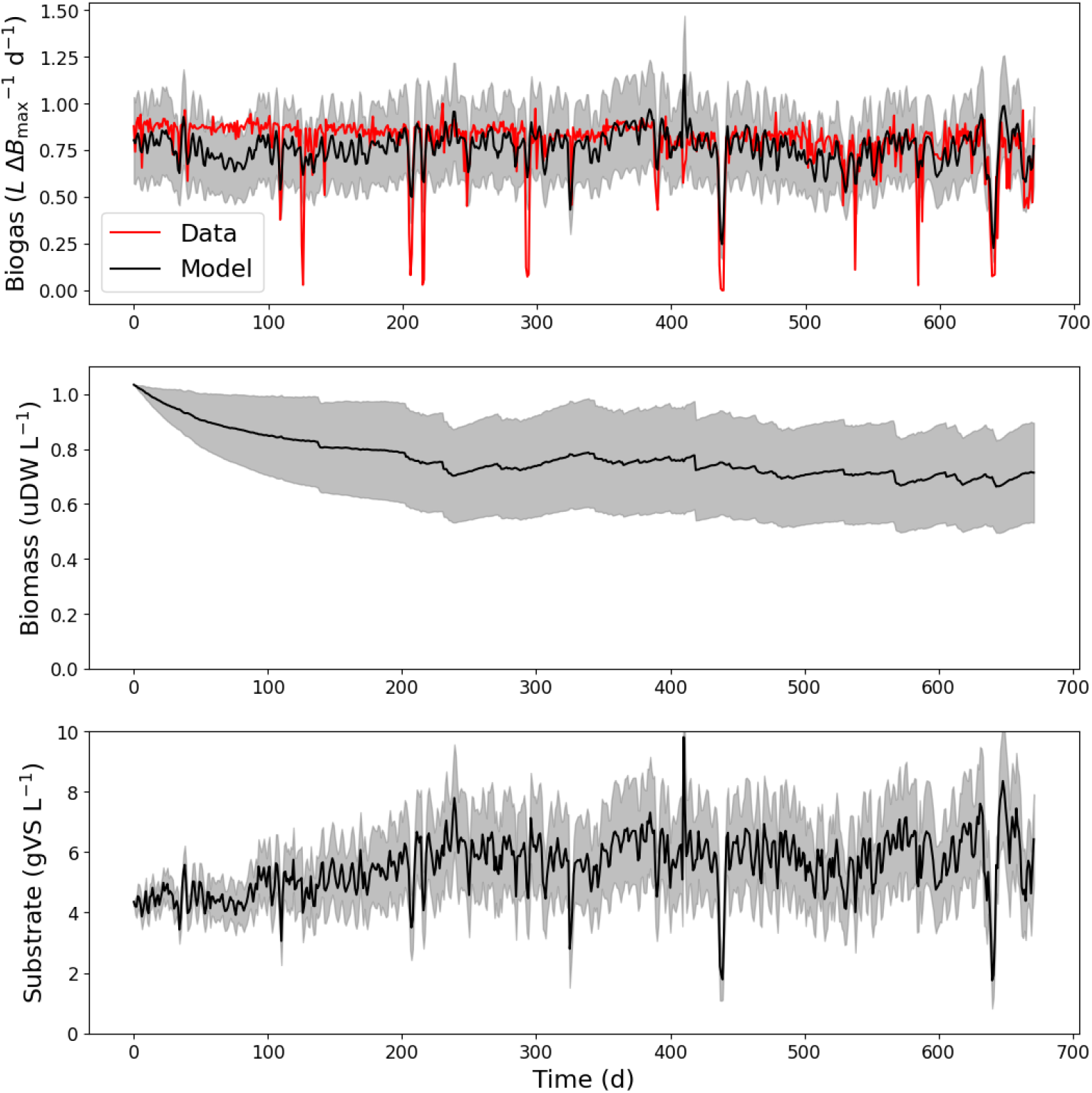
Parameter estimation fitting for site-scale digester. Gray bands are the model uncertainty (95% confidence interval)

##### Model validation

For the two lab-scale digesters, the prediction trajectory of Equation (11) yielded a MAPE of 7.5% using the parameters presented in Table 2, where cell density also maintains a steady near-unity value as shown in Figure 6. The strong performance on validation confirms the applicability of the model, particularly to the lab-scale digester.

**Figure 6.**
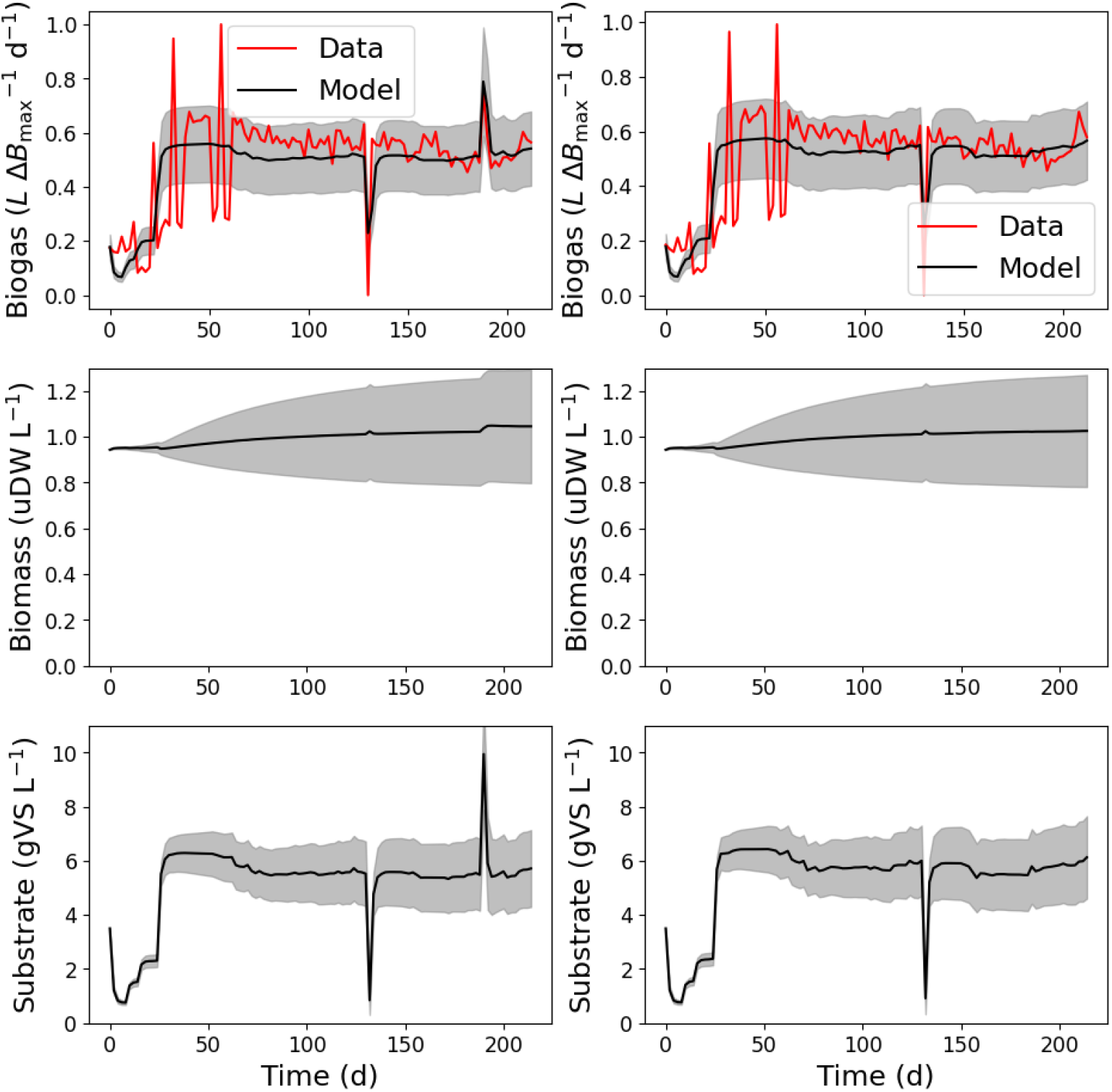
Validation fitting for lab-scale digesters 3 (left) and 4 (right). Gray bands are the model uncertainty (95% confidence interval)

#### 4.2.2 Nominal operation trajectory identification

Since *OLR*(*t*) = *S*_in_(*t*) · *HLR*(*t*), the dynamic model of the site-scale digester can be written in the compact form as in Equation (1) by assuming a known constant inflow concentration (*S*_in,fixed_). In other words, *S*_in_(*t*) = *S*_in,fixed_ = *θ*, with the state vector 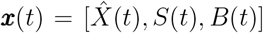, the control input *u*(*t*) = *HLR*(*t*), and the uncertain model parameter vector 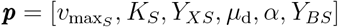. To establish the operational space, we first select the nominal operating trajectory of the control input, *HLR*, after consulting with our AD industrial partners. Specifically, the optimized nominal trajectory is obtained by balancing multiple optimization goals. These include maximizing biogas production to enhance overall output, maximizing net profit by considering both operational costs and profitability, and minimizing environmental impact by reducing the GWP associated with the process. We refer readers to [46] for detailed explanation of the multi-objective problem formulation and its solution method. This multi-objective approach ensures that the digester operates efficiently and sustainably, obtaining a balance between economic viability and environmental responsibility while maintaining high levels of methane yield. The current nominal site operation for a time horizon of 60 days is shown in Figure 7a, with the optimized nominal control trajectory depicted in Figure 7b. Here, *HLR* remains fixed throughout the day but can be adjusted on a daily basis. Both the current and optimized control trajectories suggest that it is important to alter between high and low *HLR* during operation in order to balance different process objectives.

**Figure 7.**
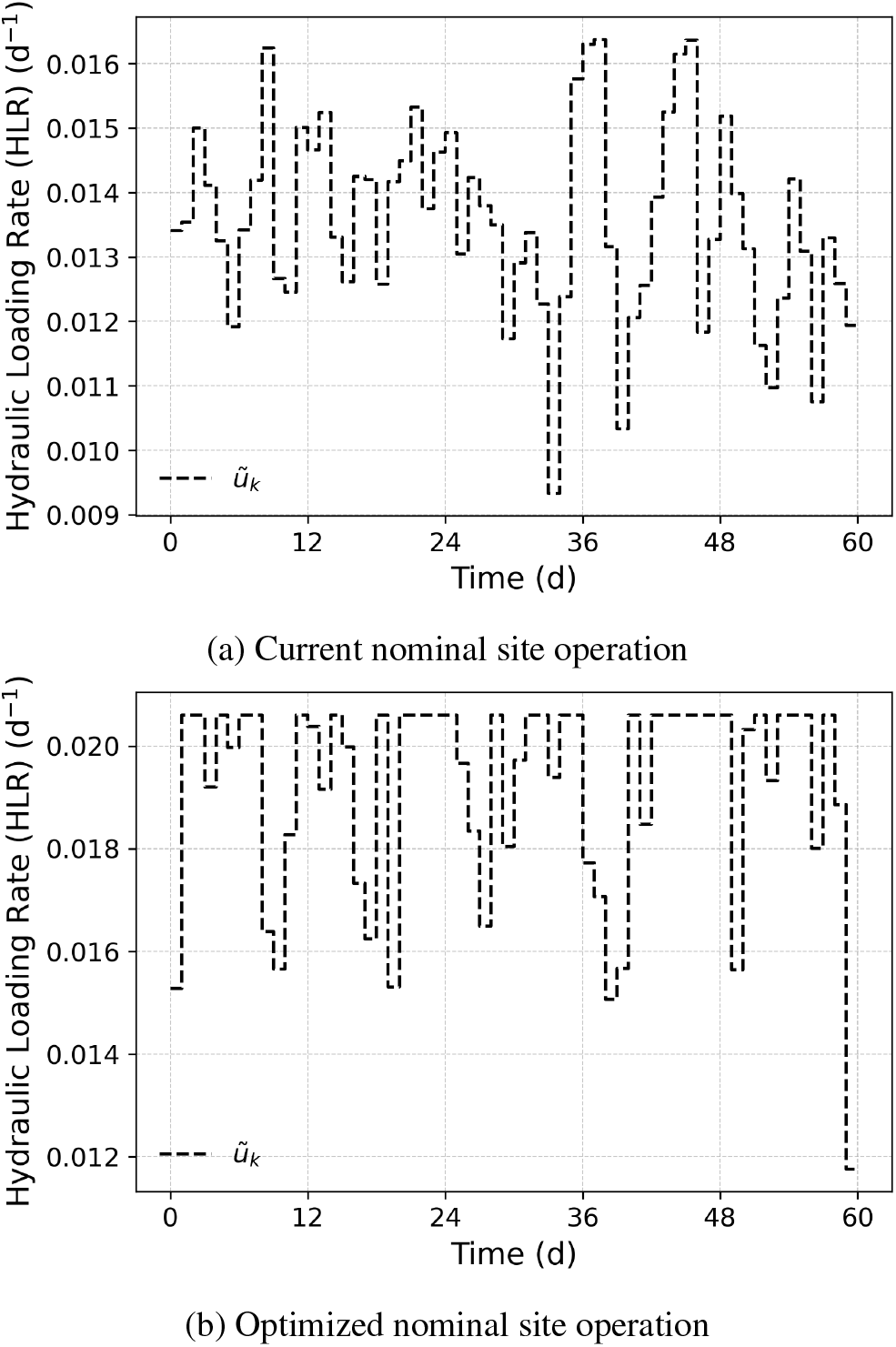
Comparison of the current nominal (a) and the optimized nominal (b) control trajectory for the site-scale digester

#### 4.2.3 Operational space design

The considered uncertainty ranges for the parameters are shown in Table 2, which intend to cover the entire experimental data trajectories observed across different digesters. These uncertainty ranges aim to capture variations in experimental responses that are likely attributed to model-process mismatches arising from parameter variability. On the other hand, sharp changes caused by disruptions such as power cut-off (see, for example, at around 25 days for lab-scale digester 1 and 2 in Figure 4) are out of the scope of this study and were not considered.

The KPI in this case is to ensure the resulting biogas production at the end of the operation period is higher than a threshold value suggested by the industry. Several path constraints are involved: (*i*) all the state and control variables are non-negative; (*ii*) rate of change of each adjacent hydraulic retention time (*HRT*, which equals to 1*/HLR*) should not exceed 35 days; (*iii*) *HRT* has to be at least 45 days; (*iv*) *HLR* should not exceed the equilibrium value to avoid wash out. The path constraints and KPI criteria are first integrated into the optimization problem (4) to determine the worst-case parameter combination by minimizing biogas production, i.e., setting the objective function *J* (***p***) to -*B*(*N* ). Subsequently, using the identified ***p***_wc_, Problems (5) and (6) are solved to determine the operational bounds while ensuring compliance with the KPI requirements and all path constraints.

#### 4.2.4. Results and discussions

The worst-case parameter combinations, ***p***_wc_, was identified by solving (4), which took 1247.7 seconds of CPU time, and was found to be [4.52 × 10^5^, 5.18 × 10^5^, 2.31 × 10^−3^, 5.38 × 10^−6^, 1.26, 1.54]. We compare the performance of two control trajectories: the optimized nominal control (NC) and the current nominal control (C). Each control trajectory is evaluated using two sets of model parameters: the nominal model parameters (NP) and the worst-case model parameters (WC). We refer to these combinations as NC-NP, NC-WC, C-NP, and C-WC in subsequent discussions. The KPI, which quantifies the final normalized biogas volume (in unit of 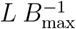, with *B*_max_ as the normalization factor) is assessed against a compliance target of 0.92. Under the optimized nominal control, the KPI is 1.19 with nominal parameters (NC-NP) and 0.97 with worst-case parameters (NC-WC); both values exceed the target. In contrast, the current nominal control achieves KPI values of 0.85 under nominal parameters (C-NP) and 0.69 under worst-case parameters (C-WC), both falling below the compliance threshold. Notably, the KPI derived from actual site data (0.86) closely matches the model’s prediction (0.85) for the current nominal control with nominal parameters.

Figure 8 illustrates the corresponding state profiles for biogas production across these four scenarios (NC-NP, NC-WC, C-NP, and C-WC), and includes experimental data for the current control with nominal parameters, i.e., C-NP. The normalized biogas production rate (top row) reveals significant differences between KPI compliant (NC-NP and NC-WC) and non-compliant (C-NP and C-WC) trajectories, with production rates under worst-case parameters consistently lower than those under nominal parameters. Biomass profiles (middle row) remain relatively consistent across all conditions, exhibiting only minor declines in WC scenarios. Substrate dynamics (bottom row), however, show more noticeable variations depending on the control strategy and parameter pairing. These findings highlight the effectiveness of optimized nominal controls in achieving compliance compared to current site operations. Nevertheless, both current and optimized nominal controls result in stable operations with nominal and worst-case parameters.

**Figure 8.**
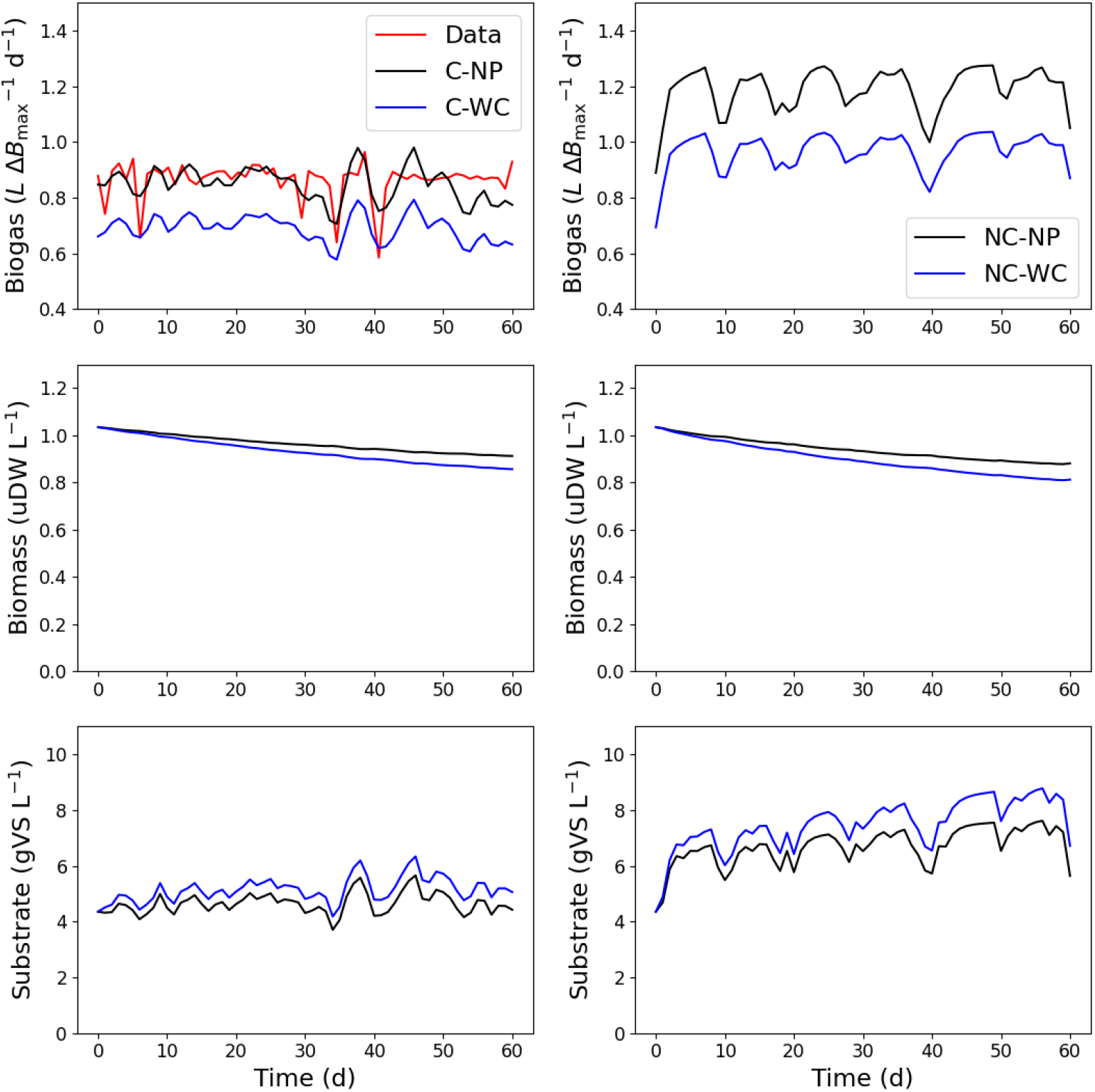
State profiles for biogas production under current nominal control trajectories (C, left subplots) and optimized nominal control trajectories (NC, right subplots), paired with either nominal (NP) or worst-case (WC) parameters.

The operational space of *HLR* around the optimal nominal control trajectory is determined from (5) and (6), which in total took 65.31 seconds of CPU time. The resulting operational space is shown in Figure 9 (shaded), allowing deviations of up to 15% from the nominal trajectory. This operational space was validated using MC simulation, as detailed in Section 3.2.4, with the maximum allowable violation rate *ξ* set at 2%. As in the first case study, 10,000 random uncertain model parameter combinations are sampled to span the entire operational space, with each sampled combination paired with a randomly generated control trajectory from the identified operational space. Figure 10 presents all the sampled uncertain model parameter combinations and their associated KPI values in a parallel coordinate plot, clearly illustrating the comprehensive exploration of parameter variability during the validation process. In the plot, the left-most vertical axis indicates KPI satisfaction, specifically whether the final normalized biogas volume exceeds the threshold of 0.92. The subsequent vertical axes represent individual parameters and their uncertain ranges, with the connecting lines illustrating the combinations of uncertain model parameters and the resulting normalized biogas production for each scenario. Notably, across all 10,000 sampled scenarios, no violations of the KPI threshold were observed.

**Figure 9.**
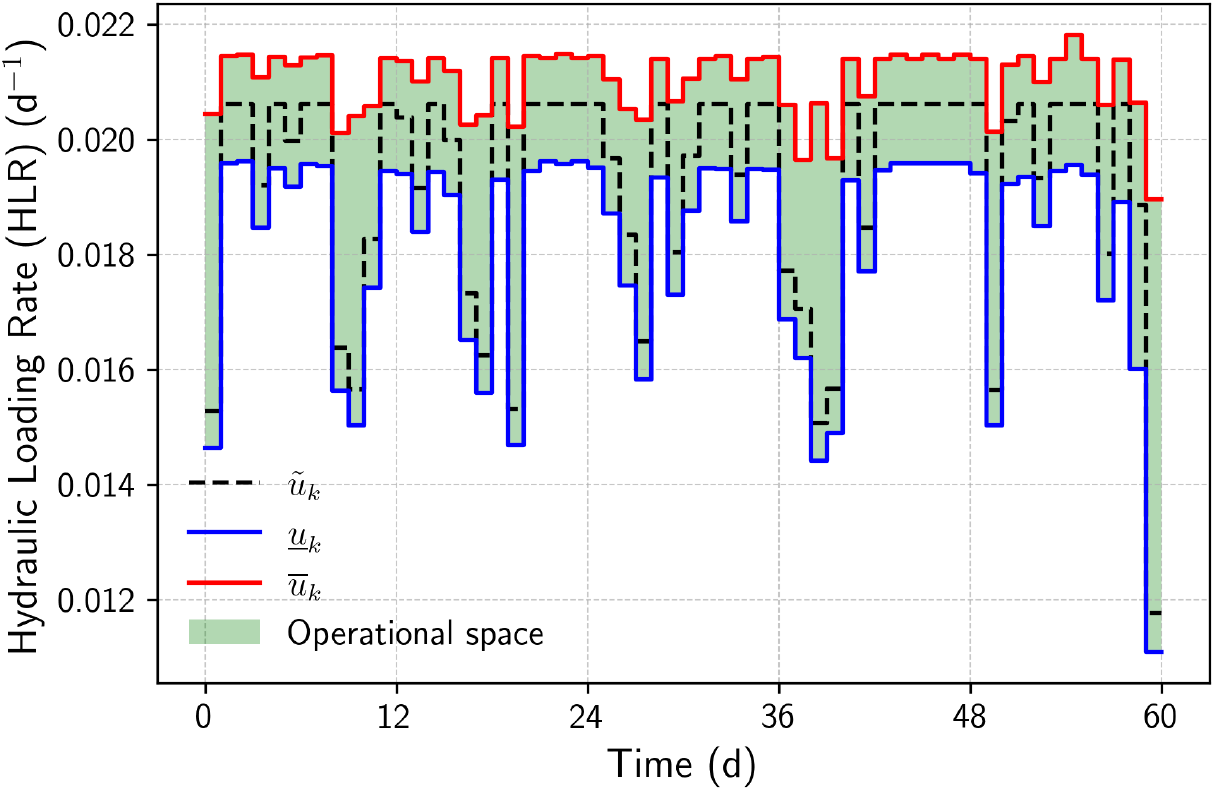
Designed operational space (shaded) for biogas production using the proposed method

**Figure 10.**
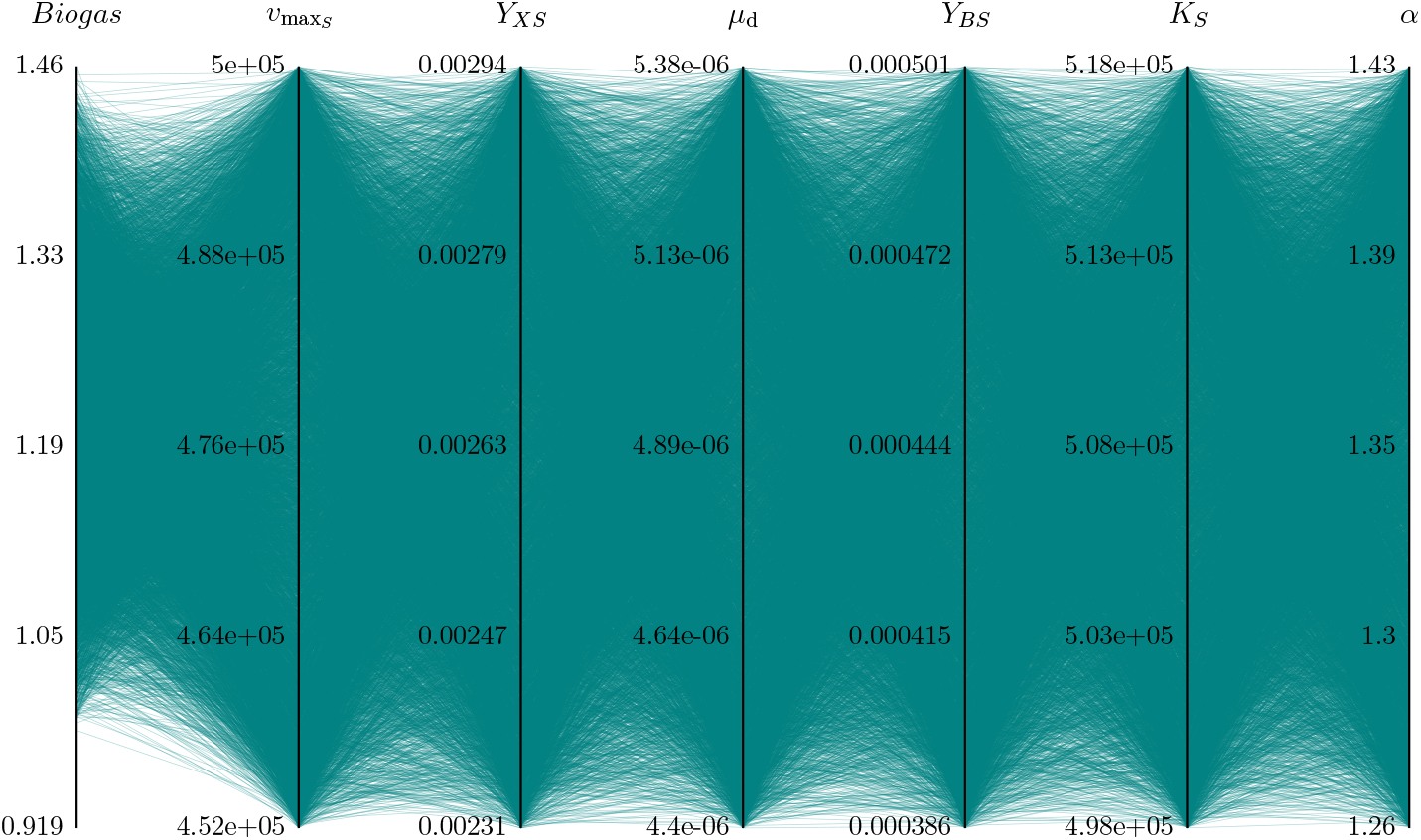
Parallel coordinate plot validating the operational space for biogas production using 10,000 scenarios. The left-most vertical bar represents KPI satisfaction (final normalized biogas volume ≥ 0.92), while the remaining bars show combinations of uncertain model parameters sampled in the MC simulation

Figure 11 presents state deviation profiles for biogas production across 10,000 scenarios under three conditions: RC-RP (random control trajectories coupled with random uncertain model parameter combinations), RC-NP (random control trajectories coupled with nominal uncertain model parameter combination), and RC-WC (random control trajectories coupled with worst-case uncertain model parameter combination). Shaded areas indicate deviation ranges within each condition, while solid and dashed lines show mean trajectories. The biogas production rate (top plot) varies noticeably, with RC-RP exhibiting the widest deviation range due to fully random parameter sampling. RC-NP and RC-WC show narrower ranges, with worst-case parameters consistently yielding lower biogas compared to nominal parameters. Biomass (middle plot) and substrate (bottom plot) profiles remain stable across conditions, showing minor deviations for RC-NP and RC-WC but larger variability for RC-RP. Despite these variations, no system constraint violations occurred, highlighting the control strategy’s reliability across diverse conditions and ensuring stable operations.

**Figure 11.**
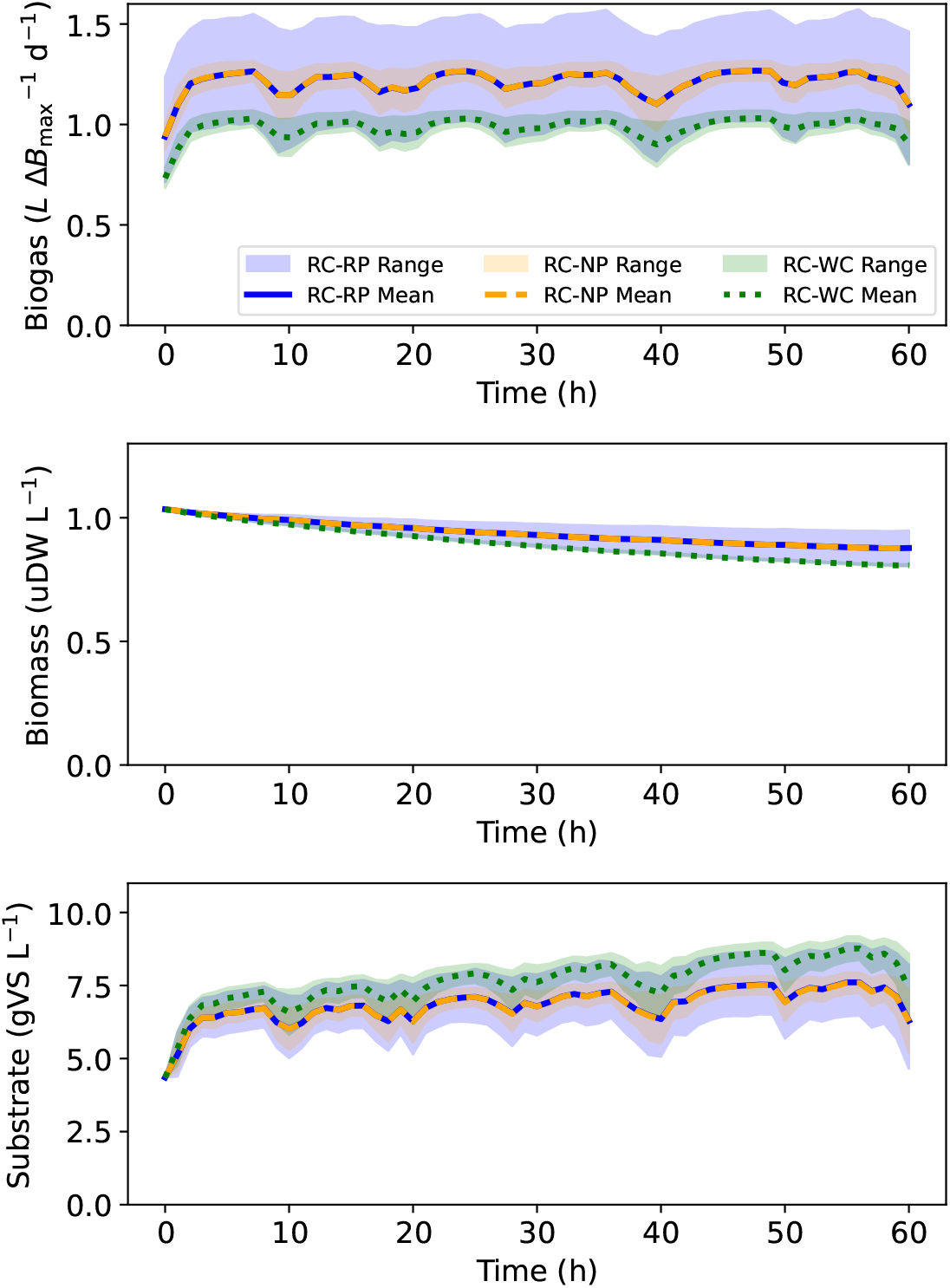
State deviation profiles for biogas production across 10,000 scenarios under RC-RP, RC-NP, and RC-WC conditions. “RC” represents randomly sampled control trajectories, with “RP”, ‘NP”, and “WC” represent randomly sampled, nominal, and worst-case parameter combinations, respectively. Lines indicate the mean, and shaded areas show the deviation range. All random samples are drawn from uniform distributions within their respective bounds

Figure 12 shows distributions of normalized KPI outcomes from 10,000 scenarios under various conditions. These include RC-RP, RC-NP, and RC-WC, as well as two additional cases: C-RP (current nominal control with random parameter combinations) and NC-RP (optimized nominal control with random parameter combinations). The dashed line at 0.92 represents the KPI target. Among the considered conditions, RC-RP, RC-NP, RC-WC, and NC-RP consistently achieve KPI outcomes above the target without any violations. In contrast, the C-RP condition exhibits a high violation rate (82.12%), highlighting the limitations of the current nominal control when combined with random parameters. The inset provides a closer look at the overlapping distributions of RC-RP and NC-RP, which exhibit a wide range of variability, while RC-NP and RC-WC show narrower variances. These results suggest that model parameter uncertainty has a greater impact on KPI variability than random control trajectories. Overall, the figure illustrates the effectiveness of the identified operational space in meeting KPI targets under uncertain conditions.

**Figure 12.**
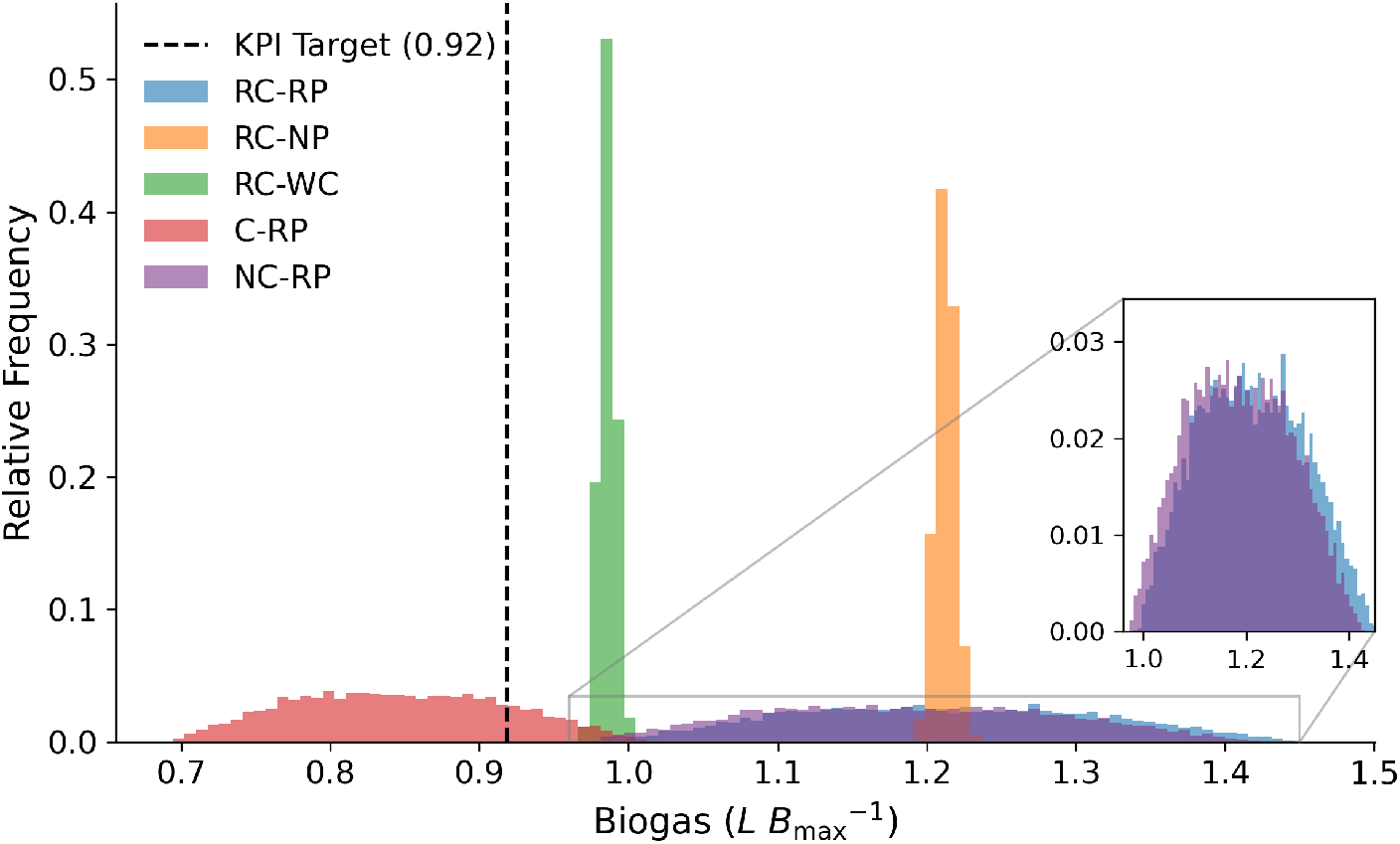
Histogram of KPI outcomes (final biogas production) across 10,000 scenarios under different conditions. The first symbol denotes the control trajectory: randomly sampled (RC), current nominal (C), or optimized nominal (NC), while the second denotes the parameter combination: randomly sampled (RP), nominal (NP), or worst-case (WC). %Violation_RC−RP_ = %Violation_RC−NP_ = %Violation_RC−WC_ = %Violation_NC−RP_ = 0, and %Violation_C−RP_ = 82.12% > *ξ* (violation threshold *ξ* = 2%). All random samples are drawn from uniform distributions within their respective bounds

## 5 Conclusion and future work

This study proposed a novel methodological framework to address uncertainties inherent in bioprocesses by introducing the worst-case operational space design (WC-OSD) approach. By focusing on designing operational spaces that pre-emptively account for uncertainties, the method ensures performance objectives are achieved even under adverse conditions. The framework was validated through two case studies: a lab-scale fermentation process for astaxanthin production and a site-scale anaerobic digestion system for biogas production. The results demonstrated the effectiveness of the proposed approach in maintaining KPIs under various uncertainty scenarios. Furthermore, the integration of symbolic optimization with scenario-based validation enhanced the reliability of the operational space, enabling scalable bioprocess solutions and reducing the need for extensive real-time monitoring infrastructure.

While the findings highlight the effectiveness of the proposed method, there remains scope for further refinement and exploration. One key assumption in this study—that the KPI can still be met under the identified nominal control trajectory despite worst-case parameter variations (as defined in Problem (4))—may not always hold in practice. In many real-world scenarios, the control trajectory that optimizes production or other objective functions under nominal conditions may fail to satisfy performance requirements under worst-case conditions. To enhance the robustness of this approach, future work could explore relaxing the KPI constraints in Problem (4) and incorporating an additional step after worst-case parameter identification to determine a new nominal control trajectory that explicitly ensures KPI satisfaction under worst-case conditions. While this adjustment may not yield maximum production or optimize other objective functions under nominal conditions, it would improve the practical applicability and reliability of the framework in real-world bioprocess operations.

Beyond addressing this assumption, further enhancements could be made to improve the framework’s adaptability to more complex and variable operational environments. For instance, incorporating additional sources of uncertainty, such as feedstock variation, would extend its applicability to dynamic bioprocess conditions. Furthermore, applying the WCOSD approach to other biochemical systems, such as pharmaceutical manufacturing or wastewater treatment, could help assess its versatility and scalability across industries.

## Acknowledgments

This work is supported by the Engineering and Physical Sciences Research Council [grant number EP/Y005600/1]; and the Biotechnology and Biological Sciences Research Council’s Doctoral Training Partnership. The authors would like to acknowledge Future Biogas Ltd. for providing support for the research in this paper.

